# Neuronal adaptation reveals a suboptimal decoding of orientation tuned populations in the mouse visual cortex

**DOI:** 10.1101/433722

**Authors:** Miaomiao Jin, Jeffrey M. Beck, Lindsey L. Glickfeld

## Abstract

Sensory information is encoded by populations of cortical neurons. Yet, it is unknown how this information is used for even simple perceptual choices such as discriminating orientation. To determine the computation underlying this perceptual choice, we took advantage of the robust adaptation in the mouse visual system. We find that adaptation increases animals’ thresholds for orientation discrimination. This was unexpected since optimal computations that take advantage of all available sensory information predict that the shift in tuning and increase in signal-to-noise ratio in the adapted condition should improve discrimination. Instead, we find that the effects of adaptation on behavior can be explained by the appropriate reliance of the perceptual choice circuits on target preferring neurons, but the failure to discount neurons that prefer the distractor. This suggests that to solve this task the circuit has adopted a suboptimal strategy that discards important task-related information to implement a feed-forward visual computation.

## Introduction

Sensory processing supports the transformation of signals from the outside world into a neural code represented by the spiking activity of cortical neurons (Hubel and Wiesel, 1959; Dubner and Zeki, 1971; Desimone et al., 1984). Decades of causal and correlative studies suggest that these representations are the basis for perceptual choice (Salzman et al., 1990; Schiller, 1993; Britten et al., 1996; Hung et al., 2005). However, it is still not understood how these sensory representations are actually combined, and what information is used, to compute a perceptual choice.

Here we focus on understanding how decision-making circuits compute perceptual choices about the orientation of a visual stimulus. This is a quintessential computation that relies on the representations encoded in the primary visual cortex (V1) (Glickfeld et al., 2013; Petruno et al., 2013; Poort et al., 2015; Resulaj et al., 2018). Perhaps the most well-established model for how such a choice is made requires that it is implemented by a neural circuit capable of monitoring the entirety of a heterogeneously tuned cortical population to estimate absolute stimulus values (Georgopoulos et al., 1986; Pouget et al., 2003; Jazayeri and Movshon, 2006; Ma et al., 2006; Graf et al., 2011). This orientation identification strategy is attractive in that, once it is learned, the same computation can be used to generalize across multiple tasks (i.e. detection and discrimination). However, this computation likely requires complex circuits (with knowledge of the full tuning curve of each neuron) and learning rules to act upon the combined output of a diversely tuned population (Deneve et al., 1999).

Instead, when faced with perceptual choices, human and animal subjects often implement task-specific strategies that require less complex circuits and learning rules (Zhang et al., 2010; Fulvio et al., 2014; Yu et al., 2017; Djurdjevic et al., 2018). Such a task-specific computation might directly evaluate the identity or level of activity within distinct neuronal ensembles, agnostic to the tuning of neurons in those ensembles, as in a linear classifier. By removing the need for an absolute stimulus estimate, the circuits that compute perceptual choice may operate faster and be more amenable to simple cellular associative learning rules.

Our goal is to understand how these computational approaches are realized by biological circuits, and how sensory information is integrated by these circuits to make perceptual decisions. Stimulus-specific adaptation is a useful tool for evaluating how sensory information is used to guide perceptual choice since it has predictable effects on neuronal activity and sensory encoding (Müller et al., 1999; Dragoi et al., 2000). By sparsifying and increasing the signal-to-noise of neuronal population responses, stimulus-specific adaptation is expected to increase the information about the presented orientation (Ulanovsky et al., 2003; Wark et al., 2007). In addition, if the stimulus orientation is estimated, adaptation to the distractor is also expected to improve orientation discrimination thresholds by decreasing the contribution of the distractor-preferring neurons (Müller et al., 1999; Dragoi et al., 2000; Kohn and Movshon, 2004; Stocker and Simoncelli, 2006). Indeed, common perceptual illusions such as the tilt after-effect and the waterfall illusion are consistent with such repulsive effects of adaptation on tuned populations (Levinson and Sekuler, 1976; Clifford, 2002; Zavitz et al., 2016). While most effects of adaptation indicate that it will improve discrimination thresholds (Kohn, 2007), there are also examples of adaptation impairing discrimination (Regan and Beverley, 1985; Ollerenshaw et al., 2014). This could be due to differences in the effects of adaptation on task-related information encoded in the cortex, or how that information is used by downstream circuits to enable behavior.

We find that adaptation in mouse visual cortex increases orientation discrimination thresholds. This is surprising since the effects of adaptation that we observe on visually responsive neurons increase the information about orientation in V1. This inconsistency between behavioral and neural results suggests that the animal fails to make use of the extra information present in the adapted population response. Indeed, both a neural decoder fit to the behavioral data and additional psychophysical experiments suggest that the animal is relying primarily on neurons that are tuned to the target stimuli. Thus, in this behavior, the underlying perceptual choice circuit does not utilize either a robust identification of stimulus orientation or an optimal task-specific computation. The utilization of only the minimal necessary information, despite costs to performance, may be the result of a prioritization of rapid processing and simple learning rules.

## Results

### Adaptation has prolonged effects on the amplitude and selectivity of visual responses

In order to understand how adaptation impacts sensory encoding, and therefore influences performance on an orientation discrimination task, we first sought to characterize the time course of adaptation in the mouse primary visual cortex (V1). Using video-rate two-photon imaging, we measured visually-evoked responses in layer 2/3 of V1 in alert mice transgenically expressing the calcium indicator GCaMP6 (see Methods). Mice passively viewed pairs of brief, identical static gratings (100 ms) presented at a range of inter-stimulus-intervals (ISIs: 0.25–4 s; **Figure 1a**). At short intervals, neurons in V1 have significantly reduced responses to the second stimulus and gradually recover (tau=592 ms) with increasing ISI (n=245 cells, 5 mice; one-way anova (p<10^−17^) with post-hoc Tukey HSD compared to non-adapted responses (250 ms: p<10^−7^; 500 ms: p<10^−7^; 1 s: p<0.0001; 2 s: p=1.0; 4 s: p=0.96); **Figure 1b-c**). We do not think that this strong adaptation is an artifact of either indicator or spike rate saturation because there was no relationship between response amplitude and degree of adaptation either within (normalized dF/F after 250 ms ISI for preferred versus neighboring orientation: p=0.37; Wilcoxon rank sum test; **Figure 1d**) or across cells (linear regression: r^2^=0.003, p=0.42; **Figure 1e**). Further, the degree of adaptation measured with extracellular single unit recording was similar to, though significantly stronger than, data collected with calcium imaging (two-way anova: main effect of recording method: p<0.001; **Figure 1f-h**). Thus, the effects of adaptation are strong and relatively long-lasting compared to the duration of the stimulus.

**Figure 1.**
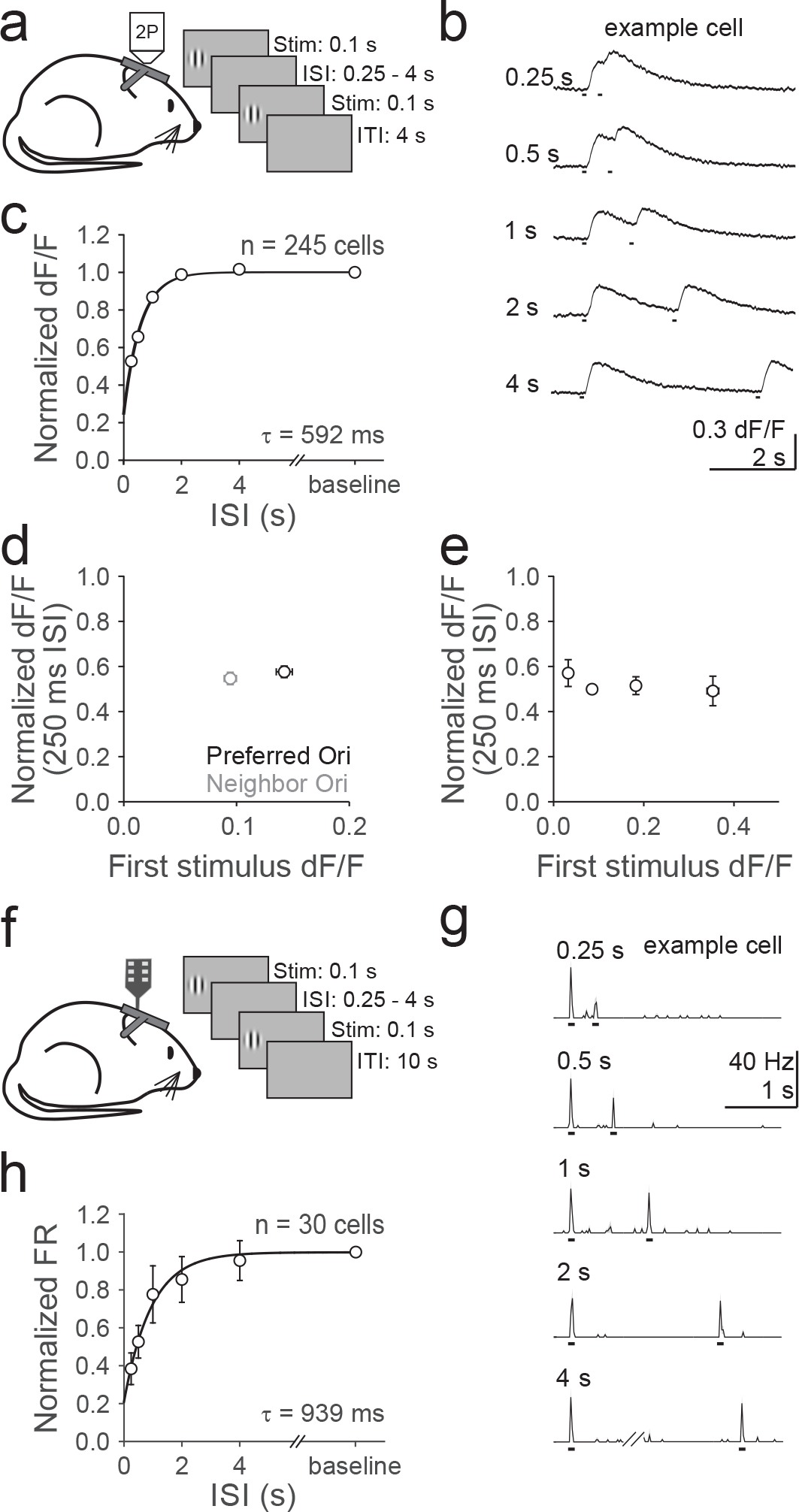
Layer 2/3 neurons in mouse V1 undergo strong adaptation. (**a**) Schematic of in vivo two-photon calcium imaging and visual stimulus protocol. Head-restrained mice passively view presentation of pairs of iso-oriented stimuli. ISI- inter-stimulus interval; ITI-inter-trial interval. (**b**) Average fluorescence traces (dF/F) from an example neuron to pairs of iso-oriented gratings separated by increasing ISIs (top to bottom). (**c**) Summary of the average amplitude of the second stimulus normalized to the amplitude of the first stimulus for each ISI for all cells (n = 245, 5 mice). Data were fit with a single exponential decay with τ = 592 ms. (**d**) Average normalized dF/F (250 ms ISI) and average dF/F in response to the first stimulus for preferred (black) and neighboring (gray) orientations. (**e**) Average normalized dF/F (250 ms ISI) for cells binned by their response to the first stimulus. (**f-h**) Same as **a-c**, for in vivo extracellular recording (n=30 cells, 4 mice). FR-firing rate. Error bars are SEM across cells.

A large component of cortical adaptation is stimulus-specific, and can thus have diverse effects that depend on the difference between a neuron’s preferred orientation and the adapter (Müller et al., 1999; Dragoi et al., 2000; Stroud et al., 2012; Patterson et al., 2013). To determine how adaptation alters orientation tuning in mouse V1, and predict how these effects might impact discrimination, we measured the orientation tuning of a population of layer 2/3 neurons with and without adaptation to a vertical grating (**Figure 2a-c**). Adaptation significantly reduces responses to stimuli near the adapter orientation (difference in normalized dF/F: two-way anova, main effect of orientation – p<10^−6^; n=241 cells, 12 mice; **Figure 2d**), and this effect is larger and affects a broader range of stimuli when the ISI is short (two-way anova, main effect of interval – p<0.0001; **Figure 2d**). Moreover, the peak responses of neurons with preferred orientations near the adapter are significantly reduced (normalized peak amplitude: two-way anova, main effect of orientation – p<10^−5^; main effect of interval – p<0.05; **Figure 2e**) and their tuning curves repelled away from the adapter (change in preferred orientation: two-way anova, main effect of orientation – p<10^−24^; main effect of interval – p<0.01; **Figure 2f**). In addition, neurons with preferred orientations orthogonal to the adapter have a significant increase in orientation selectivity index (OSI; difference from control OSI: two-way anova, main effect of orientation – p<0.01; main effect of interval – p<0.01; **Figure 2g**), likely due to selective adaptation of responses on the flanks of their tuning curves (Dragoi et al., 2002). Thus, adaptation alters the amplitude, preference and selectivity of neuronal responses in V1 in a manner very similar to what has been previously observed in carnivores and primates (Müller et al., 1999; Dragoi et al., 2000; Patterson et al., 2013).

**Figure 2.**
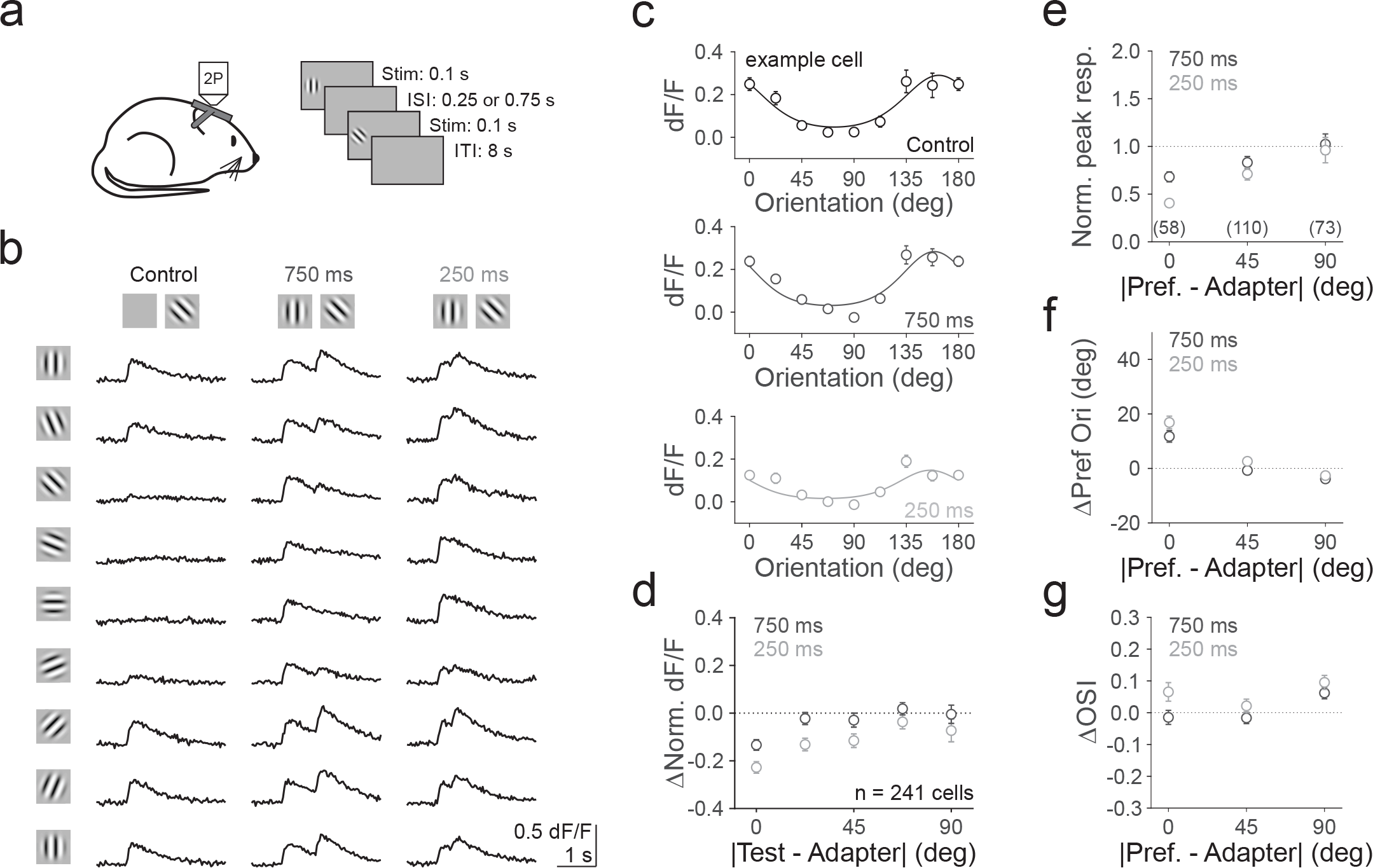
Adaptation changes the orientation tuning and preference of layer 2/3 neurons in mouse V1. (**a**) Schematic of in vivo two-photon calcium imaging and visual stimulus protocol. Head-restrained mice passively view presentation of pairs of stimuli with varying orientations and intervals. (**b**) Average dF/F from an example cell to eight different orientations (rows) without adaptation (left column: Control) or after 750 (middle column) or 250 ms (right column) recovery from adaptation to a vertical (0°) grating. (**c**) Average orientation tuning curves for the neuron in **b** measured in control (top, black) and after 750 (middle, dark gray) or 250 ms (bottom, light gray) recovery from adaptation. Average responses (error bars are SEM across trials) in each condition were fit with a von Mises function. (**d**) Summary of the average difference in dF/F (each cell normalized to its own peak response without adaptation) as a function of stimulus distance from the adapter (n = 241 cells, 12 mice) after 750 (dark gray) or 250 ms (light gray) recovery from adaptation. Error bars are SEM across cells. (**e**) Summary of the average normalized peak dF/F as a function of the distance of the cells’ preferred orientation from the adapter. Cells are binned by their preferred orientation (determined from the peak of the fit in control conditions) into three groups: less than 20°, between 20° and 70°, and more than 70° from the adapter (n = 58, 110 and 73 cells). (**f**) Summary of the average change in preferred orientation as a function of the distance of the cells’ preferred orientation from the adapter. Positive shift indicates repulsion and negative shift indicates attraction relative to the adapter. (**g**) Summary of the average change in orientation selective index (OSI) as a function of the distance of the cells’ preferred orientation from adapter. Positive change indicates increased selectivity and negative change indicates decreased selectivity relative to control.

### Orientation identification models predict that adaptation improves discrimination

Theoretical investigations using optimal decoding strategies to estimate stimulus orientation predict that adaptation to distractor stimuli should improve discrimination (Clifford et al., 2001; Stocker and Simoncelli, 2006; Zavitz et al., 2016). This is because many of these strategies work by considering the activity of each neuron as some number of votes in favor of the proposition that the presented stimulus was actually the preferred stimulus of that neuron. Adaptation thus causes a reduction in the number of votes for stimulus values near the adapting stimulus resulting in a stimulus estimate that is biased away from the adapting stimulus. More generally, biases away from the adapting stimulus can result from unbalanced changes in the signal-to-noise ratio (SNR) of the population response in which SNR increases for stimulus values near to the adapting stimulus relative to those stimulus values that are further away (Stocker and Simoncelli, 2006). The effect of the bias in stimulus estimate would be to increase both hits (correct identification of targets) and false alarms (identification of distractors as targets) by repelling the stimulus value away from the distractor. In contrast, the effect of the decrease in variance is associated with an increase in hits and a decrease in false alarms, by increasing the reliability of estimates close to the adapting stimulus.

To test these hypotheses, we applied the neural data collected from the mouse visual cortex to an optimal decoder of orientation. Specifically, for each dataset, we identified 10-15 well-tuned neurons and used their activity to empirically construct a probabilistic decoder of neural activity. The decoder assumes only that the posterior distribution across orientations given neural responses is von Mises (i.e. tuned to orientation) and that neural activity is linearly related to the log of this posterior (Ma et al., 2006). Thus, we extracted the maximum of the posterior probability distribution to reliably estimate the orientation of the stimulus presented on individual trials across all adaptation conditions (**Figure 3b**). While adaptation did not significantly alter the bias in estimated orientation, shorter ISIs significantly decreased the variability of the estimate (250 ms vs. 750 ms: p<10^−5^; F-test; **Figure 3b**) consistent with the expectation that adaptation improves the SNR in this class of models (Clifford et al., 2001; Stocker and Simoncelli, 2006; Zavitz et al., 2016).

**Figure 3.**
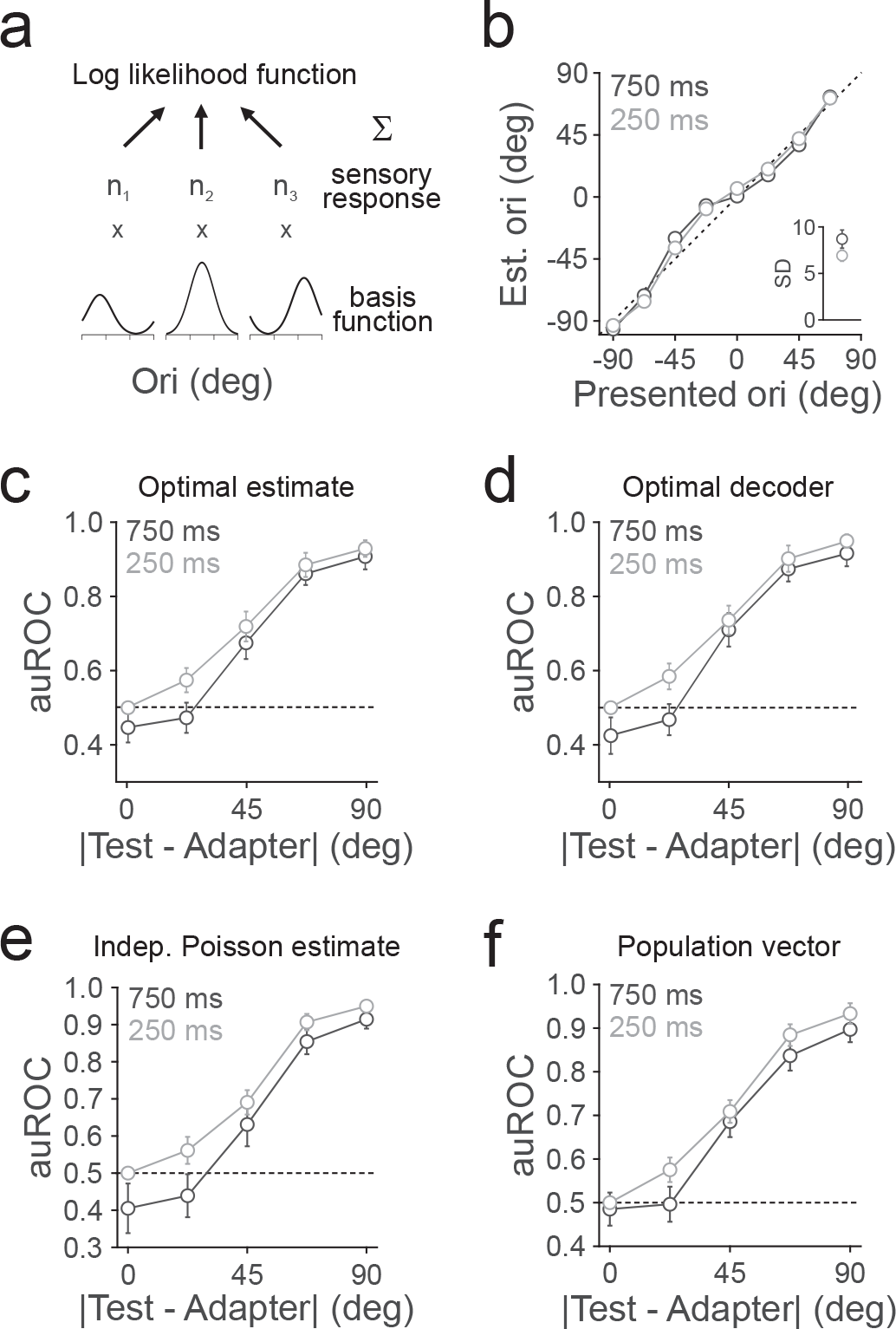
Feature identification models predict that adaptation improves discrimination. (**a**) Schematic of the orientation identification model in which the orientation of each stimulus is identified through a population code to determine its maximum likely orientation. The likelihood is determined by scaling each neuron’s basis function (see Methods) to the amplitude of its response, n_i_, to a given test stimulus and then summing across the population. The peak of this likelihood function is the estimated orientation. (**b**) Average estimate of orientation as a function of the difference of the test stimulus from the adapter after 750 (dark gray) or 250 (light gray) ms recovery from adaptation (n = 10 mice). Error bars are SEM across experiments. Inset-average standard deviation in estimate of orientation, normalized to control, as a function of adaptation state. (**c**) Summary of the average auROC, found by estimating the presented orientation, as a function of stimulus distance from the adapter after 750 or 250 ms recovery from adaptation. (**d-f**) Same as **c**, for the average posterior probability (**d**), the independent Poisson estimator (**e**), and the population vector (**f**).

In order to test the effects of adaptation on discrimination, instead of just stimulus estimation, we generated neurometric functions measuring the discriminability of the target and distractor stimuli. We compared the distributions of single trial estimates of stimulus orientation for each target stimulus to the distribution of estimates in response to the distractor stimulus (in the adapted condition to mimic the state of the distractor in our task). We summarized the average neurometric function as the area under the Receiver Operating Characteristic curve (auROC) statistic. In our data, the major effect of adaptation is to decrease the variance of the estimated orientation. Consistent with this, the auROC increased in the presence of adaptation for small orientation changes (250 ms vs. 750 ms: 22.5°: p<0.001; n=10 mice; paired t-test; **Figure 3c**) and was not significantly different for distractor stimuli (0°: p=0.14).

Similarly, alternative optimal estimators that use the posterior probability distribution as the decision variable (auROC-22.5°: p<0.0001; 0°: p=0.08 paired t-test; **Figure 3d**) or that assume independent Poisson statistics (auROC-22.5°: p<0.05; 0°: p=0.19; paired t-test; **Figure 3e**) predict an improvement in discrimination of small orientation differences. We also tried a suboptimal orientation estimator using a population vector, and again found that adaptation decreases discrimination threshold (auROC-22.5°: p<0.01; 0°: p=0.71; paired t-test; **Figure 3f)**. Thus, all variants of population-based orientation identification that we tested predict that adaptation will decrease thresholds on a discrimination task.

### Adaptation increases orientation discrimination thresholds

To determine whether adaptation in fact improves orientation discrimination, we designed a task in which the mouse needs to use information about the orientation of visual stimuli to earn reward. In this task, head-restrained mice are trained to press a lever to initiate a trial and release it to report a target orientation (**Figure 4a** and **Supplementary Movie 1**). On each trial, the lever press triggers the serial presentation of 2-9 gratings of the same orientation (“distractors”; 100 ms duration; mice were trained with either a 0° (n=9) or 45° (n=2) distractor) in which each presentation is separated by a randomly selected ISI (250, 500, or 750 ms) that prevents the mouse from anticipating the upcoming interval. The number of distractor stimuli on each trial is also variable to prevent the mouse from anticipating the target presentation (range of differences from the distractor: 9-90°). If the mouse releases the lever within a window 200-550 ms following the onset of the target stimulus, it is considered a hit; if the mouse releases the lever within the same window following a distractor stimulus, it is considered a false alarm (FA). Thus, this task allows us to compare the discrimination threshold and FA rate for stimuli following different ISIs, and therefore in different adaptation states.

**Figure 4.**
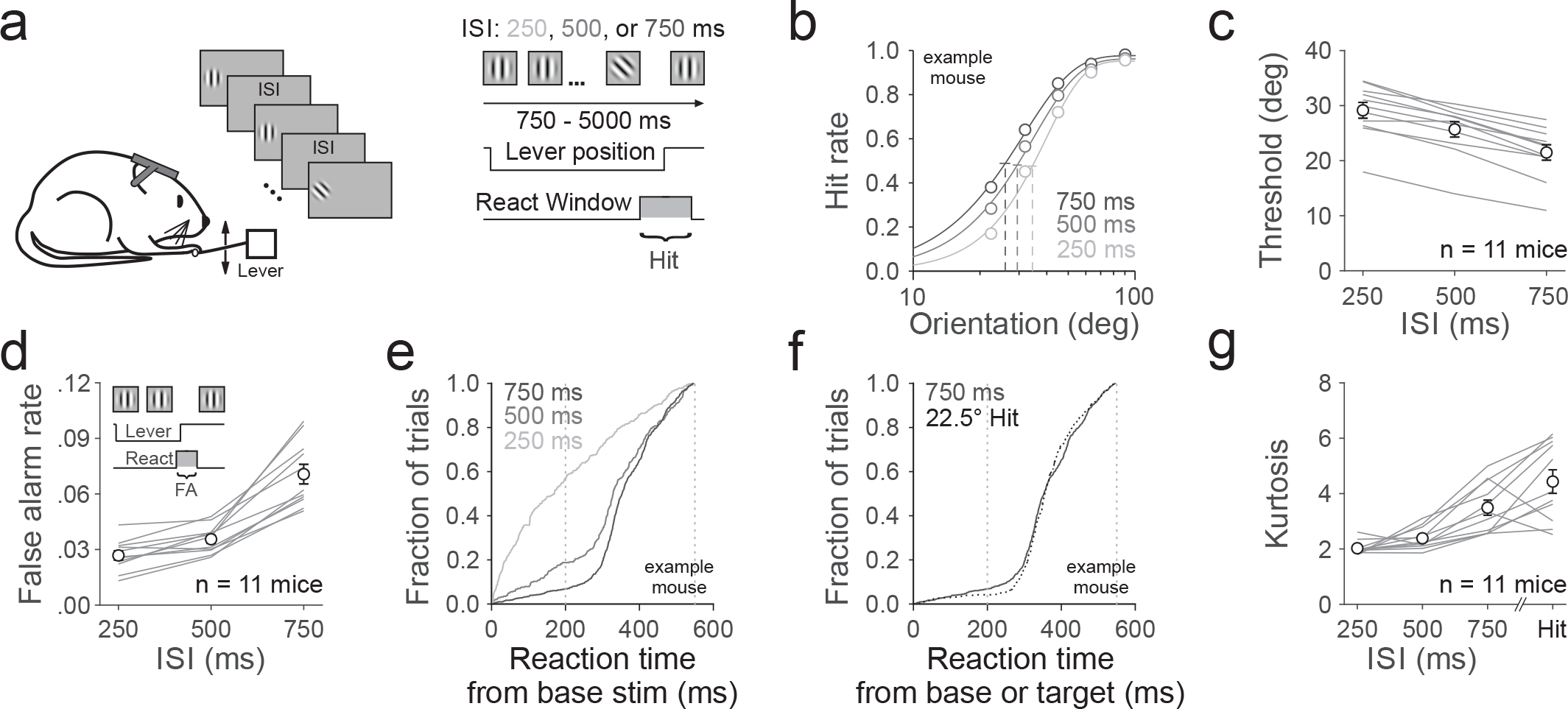
Adaptation decreases hit and false alarm rates on an orientation discrimination task. (**a**) Schematic of behavioral setup and trial progression. Head restrained mice press a lever to initiate a trial, triggering the repeated presentation of a vertically oriented grating (100 ms) with a randomly interleaved ISI (250 (light gray), 500 (gray) or 750 (dark gray) ms); the mouse must release the lever within a short window following the presentation of a non-vertical grating to receive a reward. (**b**) Hit rate for an example mouse in which target orientations were sorted according to the preceding ISI. Data are fit with a Weibull function; vertical lines denote threshold, error bars are 95% confidence intervals. (**c**) Summary of the threshold across ISIs. Open circles are the average of all mice (n = 11 mice), connected gray lines are individual mice; error bars are SEM across mice. (**d**) Summary of the average FA rate across mice. Inset: schematic illustrating the window for a FA. (**e**) Cumulative histogram of reaction times for early releases relative to the time of distractor stimulus onset sorted according to the preceding ISI; same mouse as in **b**. Vertical lines represent react window used to calculate FA rate. (**f**) Cumulative histogram of reaction times for early releases following a 750 ms interval (dark gray; same data as in **e** for comparison) or a 22.5° target (dotted line; including all ISIs). Note the similarity in the shape of the distribution. (**g**) Summary of the average kurtosis of the reaction time distributions across mice for the three intervals and the 22.5° target (including all ISIs).

We find that the discrimination threshold is increased when the ISI is short (one-way anova with post-hoc Tukey HSD compared to 250 ms: 500 ms: p=0.21; 750 ms: p=0.002; n=11 mice; **Figure 4b-c**). Although discrimination threshold decreases with trial length (p<10^−3^, one-way anova; **Figure 4 – figure supplement 1a**), and trials with a short pre-target ISI are, by definition, shorter than those with long ISIs (p<10^−13^, one-way anova; **Figure 4 – figure supplement 1c**), the ISI-dependent changes in threshold remain intact after matching the trial length across ISIs (two-way anova, main effect of method-all versus matched trials: p=0.18, **Figure 4 – figure supplement 1d**). This suggests that the effects of trial length and ISI on discrimination threshold are independent from each other. Indeed, the effect of ISI on discrimination threshold is also the same for both short and long trials (two-way anova- main effect of trial length: p=0.36; **Figure 4 – figure supplement 1e**). Importantly, we found that arousal state is stable across ISIs because there is 1) no significant difference in lapse rate across ISIs (250 ms: 0.05±0.01; 500 ms: 0.05±0.01; 750 ms: 0.05±0.01; p=0.97; one-way anova; n=11 mice); and 2) no difference in pupil size preceding correctly detected or missed targets (p=0.87, paired t-test; n=3 mice; **Figure 4 – figure supplement 2a-c**) or across ISIs (p=0.96, one-way anova; n=3 mice; **Figure 4 – figure supplement 2d-f**). We also considered the possibility that the uncertainty in the timing of stimulus appearance might be influencing the animals’ behavior, for instance by generating surprise at the appearance of stimulus earlier or later than expected. However, we found animals also have a lower threshold for longer intervals when ISIs were interleaved on a trial-by-trial rather than presentation-by-presentation basis (500 ms vs 250 ms: p=0.01, paired t-test; n=3 mice). Thus, adaptation state has an acute effect on discrimination threshold.

We also find a decrease in FA rate with adaptation (one-way anova with post-hoc Tukey HSD compared to 250 ms: 500 ms: p=0.22; 750 ms: p<10^−8^; **Figure 4d** and **Figure 4 – figure supplement 1b-d,f**). One interpretation of the observed increase in FA rate following longer intervals is that there is an increase in impatient responses during the extended ISI. If true, this would lead to an increase in release probability shortly after the stimulus on long intervals. However, inspection of the distribution of reaction times reveals the opposite effect: the distribution of releases to distractor stimuli following short intervals had shorter latencies than those following longer intervals (one-way anova with post-hoc Tukey HSD compared to 250 ms: mean reaction time: 500 ms: p<10^−6^; 750 ms: p<10^−8^; n=11 mice, example mouse in **Figure 4e**). The distributions of reaction times are consistent with there being two classes of FAs: 1) releases following distractor stimuli in which the mouse guessed that it was a target; and 2) “spontaneous” releases due to non-sensory factors (i.e. impatience). The platykurtic distribution of reaction times in the 250 ms ISI condition is a hallmark of this latter, non-sensory behavior (Tiefenau et al., 2006). In contrast, the comparatively leptokurtic distribution following the longer intervals suggests that the majority of these are stimulus-driven releases (one-way anova with post-hoc Tukey HSD compared to 250 ms: 500ms: p=0.76; 750ms: p=0.002; 22.5° target: p<10^−6^; **Figure 4g**). In fact, the distribution of responses following the 750 ms ISI closely resembled the reaction time distribution when a 22.5° target stimulus was presented (one-way anova with post-hoc Tukey HSD of 750 ms ISI distractor compared to 22.5° target: p=0.07; **Figure 4f-g**). Notably, ISI had no effect on the distribution of responses to 22.5° targets (kurtosis-250 ms: 3.8±0.3; 500 ms: 4.1±0.4; 750 ms: 3.5±0.5; p=0.62, one-way anova), suggesting that the majority of these responses are stimulus driven in all conditions. Thus, adaptation reduces the FA rate by decreasing the likelihood of a stimulus-driven response to a distractor.

In this task, adaptation decreases both hit and FA rate. Such concomitant changes in hit and FA rate are often associated with changes in bias (c) as measured using signal detection theory (Green and Swets, 1966). Indeed, adaptation does significantly increase c (22.5° target-250 ms: 1.28±0.06; 500 ms: 1.08±0.05; 750 ms: 0.75±0.06; p<10^−5^; one-way anova; n=11 mice). However, this is due to a reduction in the amplitude of both targets and distractors (**Figure 2**) thereby shifting the optimal criterion, and resulting in an increase in measured bias (Witt et al., 2015). This supports the argument that adaptation increases discrimination thresholds through its effect on sensory processing in the visual cortex.

To determine whether the activity in visual cortical circuits is necessary for the behavioral effects of adaptation on discrimination, we used an optogenetic approach to transiently suppress activity in V1 on randomly interleaved trials (**Figure 5a-c**; see Methods). The light power was titrated to decrease hit rates for small orientation differences (22.5°-p<0.05, paired t-test, n=4 mice) without affecting performance on easy trials (90°-p=0.54, paired t-test). Suppression of V1 increased the discrimination threshold for all ISIs (two-way anova, main effect of V1 inhibition - p<10^−8^; main effect of interval – p<10^−6^; **Figure 5d**) and significantly reduced the dependence of threshold on ISI (two-way anova, main effect of V1 inhibition – p<0.01; **Figure 5e**). Suppression of V1 also reduced the FA rate (two-way anova, main effect of V1 inhibition - p<10^−5^; main effect of interval – p<10^−7^; **Figure 5f**) and its dependence on ISI (two-way anova, main effect of V1 inhibition – p<0.01; **Figure 5g**). Importantly, there was no effect of either V1 inhibition or ISI on lapse rate (control-250 ms: 0.15±0.05; 500 ms: 0.12±0.05; 750 ms: 0.16±0.04; V1 inhibition-250 ms: 0.16±0.04; 500 ms: 0.16±0.04; 750 ms: 0.14±0.04; main effect of V1 inhibition: p=0.80; main effect of ISI: p=0.83; two-way anova). This suggests that circuits in V1 are involved in orientation discrimination (Glickfeld et al., 2013), and that adaptation state in these circuits impacts task performance.

**Figure 5.**
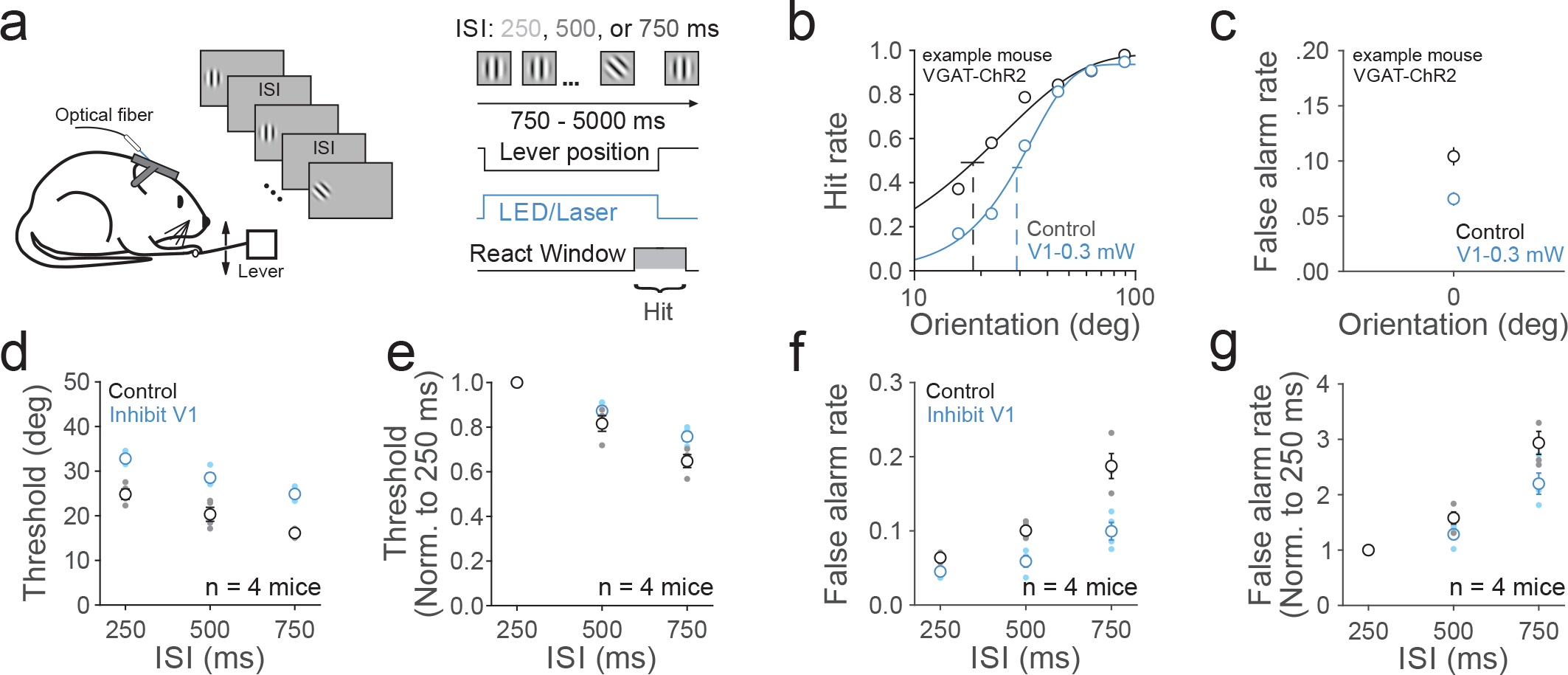
V1 is required for discriminating orientation and partially accounts for the dependence of behavior on adaptation. (**a**) Schematic of behavioral setup and trial progression. V1 inhibition was achieved via optogenetic excitation of ChR2-expressing inhibitory interneurons using VGAT-ChR2 (n = 2) and PV::Cre-AAV.flex.ChR2 (n = 2). (**b**) Hit rate on randomly interleaved control (black) and V1 inhibition (blue) trials, averaged across all ISIs, for an example VGAT-ChR2 mouse; same conventions as in Figure 4b. (**c**) Same as **b**, for FA rate. (**d**) Summary of the average threshold for control and V1 inhibition by ISI (n = 4 mice). Error bars are SEM across mice. (**e**) Summary of change in threshold, normalized to the threshold in the 250 ms condition, for each ISI. (**f-g**) Same as **d-e**, for FA rate.

### Behavioral evidence for a task-specific circuit that preferentially weights target-preferring neurons

The effects of adaptation on neuronal activity and behavior are at odds with each other. The neuronal data suggests that there is increased information about the stimulus orientation under adaptation, but the mouse clearly fails to take advantage of this information in the task. However, perception is constrained not only by the information available, but also by the computation adopted for decoding that information. Thus, our goal is to infer the perceptual choice circuit used to perform this task by identifying a computation that is consistent with the effects of adaptation on behavior: namely an increase in threshold and a decrease in FA rate.

Our previous analysis suggests that the behavior is not consistent with any of the tested computations that estimate stimulus orientation (**Figure 3**). However, there are multiple alternate solutions that the mouse could have adopted to solve this discrimination task. One such generalist strategy might be to compare each stimulus to the one that preceded it and declare whether it is the same or different (a change detection). To test whether the mouse adopts this general strategy, we perturbed the task parameters. We tested animals trained on the task in **Figure 4** on a new task where the distractor orientation could either be the trained orientation (0º) or rotated 15º or −15º from the trained orientation (randomly interleaved on a trial-by-trial basis), with the set of target orientations rotated 9-90° counter-clockwise relative to the distractor orientation (**Figure 6a**). If the mouse adopts the generalist change detection strategy, the psychometric curves for the three distractor conditions should be the same. This is because task difficulty depends only on the difference between the target and distractor orientations, not on the absolute distractor orientation (**Figure 6b**). However, we found that all six mice had lower discrimination thresholds in the 15º condition (rotated towards learned targets) when compared to the 0º condition, which in turn had lower discrimination thresholds than in the −15º condition (rotated away from learned targets; one-way anova, p<10^−4^, n=6 mice; **Figure 6d-e**). The effects on FA rates were not symmetric: all six mice had higher FA rates in the 15º condition when compared to the 0º condition (one-way anova with post-hoc Tukey HSD compared to 0º, p<0.05, n=6 mice; **Figure 6e**), but no difference in FA rate between the 0º and - 15º conditions (one-way anova with post-hoc Tukey HSD −15º compared to 0º, p=0.97). This clearly indicates that the mouse is not performing a generic change detection task.

**Figure 6.**
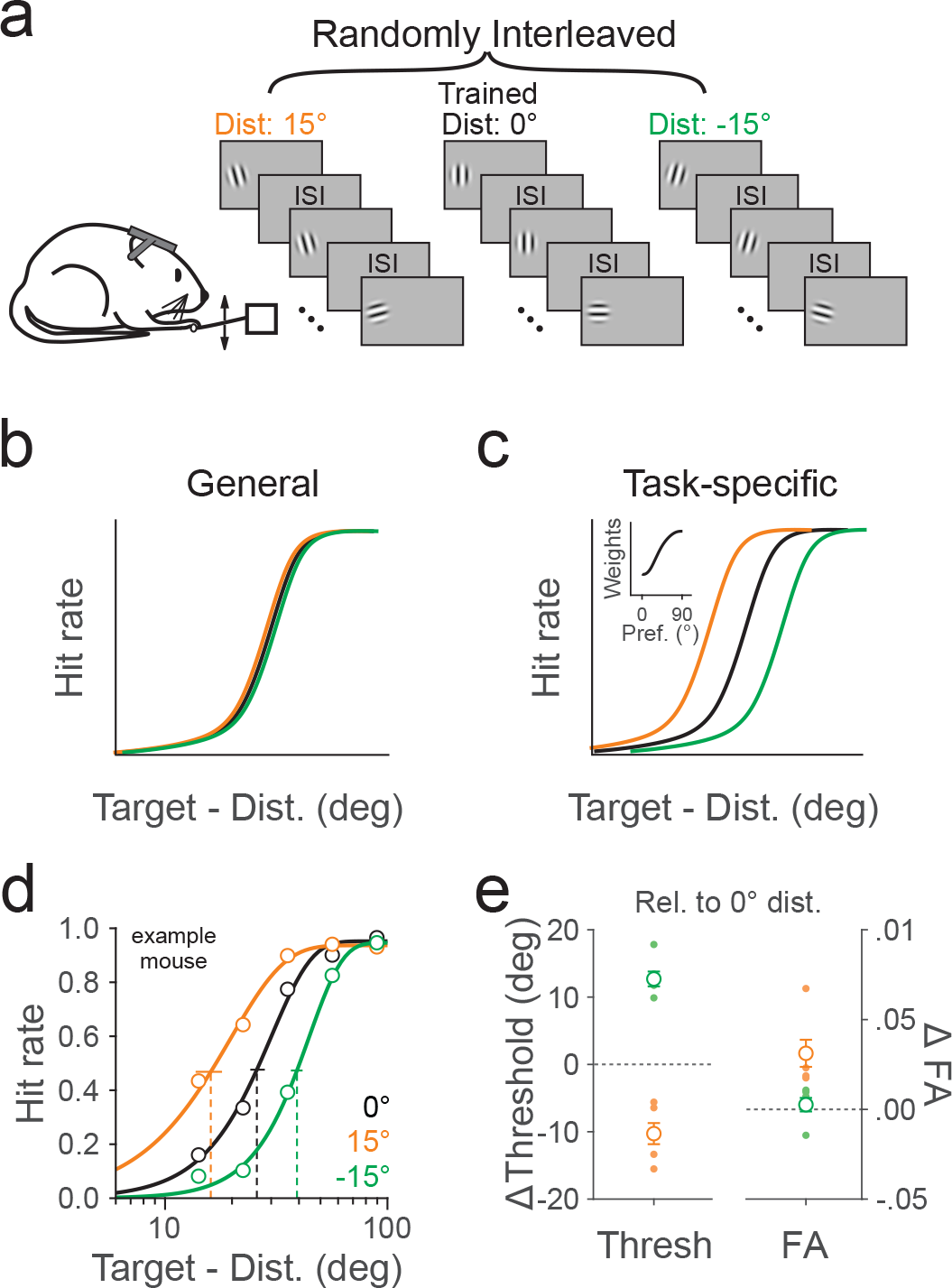
Behavior is inconsistent with a general change detection strategy, but can be explained by a task-specific circuit with biased weights. (**a**) Schematic of the trial progression. For each trial, the distractor (Dist) orientation could be 15° (orange), 0° (black) or −15° (green). The target orientations are 9° to 90° counter-clockwise from the distractor orientation. (**b**) If the mouse adopts a general change detection strategy, the discrimination threshold should be similar across different distractor orientations. (**c**) If the mouse adopts a task-specific strategy that discriminating a learned quadrant of target orientation space (positive orientations), the discrimination threshold would be lower for 15° distractor and higher for −15° distractor when compared to 0° distractor. Inset: schematic of biased weights across neurons tuned for positive orientations. (**d**) Hit rate for an example mouse in which target orientations were sorted according to the distractor orientations. (**e**) Summary of the average change in the threshold (left) and FA rate (right) for each distractor orientation relative to the 0° condition across mice. Open circles are the average of all mice (n = 6 mice), small filled circles are individual mice; error bars are SEM across mice.

Instead, the result is more consistent with a task-specific strategy where the mouse has learned that 0º gratings are distractors while all stimuli with positive orientation are targets. This can be accomplished by a perceptual choice circuit that more strongly weights neurons with preferred orientation close to 90º (**Figure 6c**).

### The effects of adaptation are consistent with a perceptual choice circuit that ignores distractor preferring neurons

The mouse’s behavior suggests that it is using a task-specific strategy that does not require orientation identification to perform this task. Thus, we hypothesized that the perceptual choice circuit might be taking advantage of a linear combination of neuronal activity that directly separates targets from distractors. To predict how the circuit might be combining the population activity to accomplish this task-specific computation, we used a linear combination of neuronal activity (from the same subpopulations of 10-15 well-fit neurons from each data set used in **Figure 3**) to fit the average behavioral data in **Figure 4**(**Figure 7 – figure supplement 1a**). Consistent with the behavioral result in **Figure 6**, the weights found from this fit were not equal across the population, with neurons preferring target orientations tending to have positive weights (orientation preference greater than 11.25° from the distractor: p<0.001; Student’s t-test **Figure 7a**). However, the weights assigned to neurons preferring stimuli closer to the distractor orientation were not significantly different from zero (orientation preference within 11.25° of the distractor: p=0.3). This was surprising since an optimal classifier should both positively weight the target orientations (to increase the probability of identifying a target stimulus) and negatively weight the distractor orientation (to decrease the probability of mistaking it for a target).

**Figure 7.**
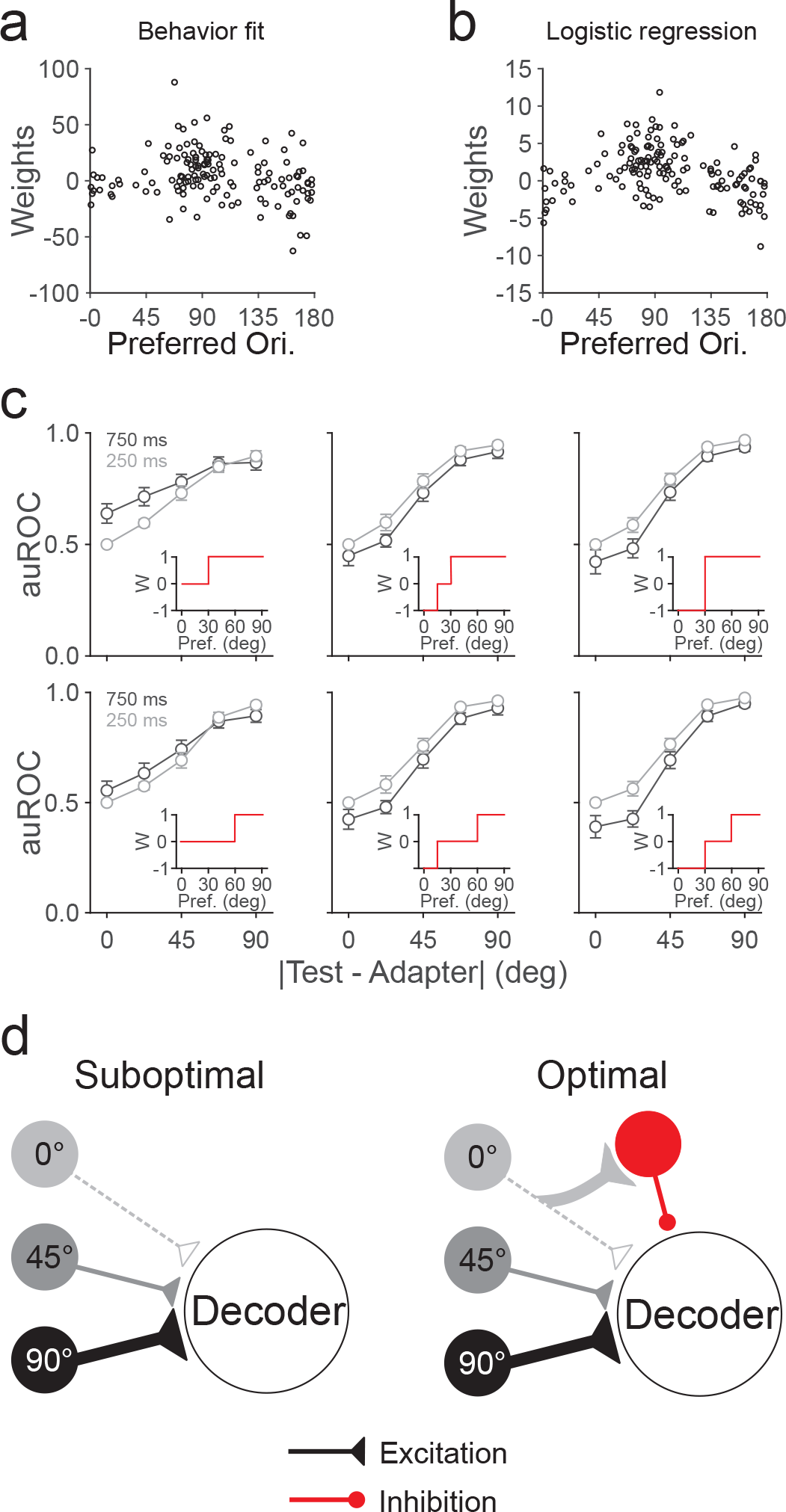
Predicted weights are biased and positive. (**a**) Summary of weights found by a fit to the behavioral data as a function of neuron orientation preference. (**b**) Same as **a**, for weights found by a logistic regression. (**c**) Summary of the average auROC as a function of stimulus distance from the adapter after 750 (dark gray) or 250 (light gray) ms recovery from adaptation, found by a weighted sum of neuronal activity (n = 10 mice). Rows: Positive weight for neurons with orientation preference greater than 30° (top) or 60° (bottom); Columns: Negative weight for neurons with orientation preferences less than 15° (middle) or 30° (right), or no negative weights (left). Insets show weighting scheme for each panel. Error bars are SEM across experiments. (**d**) Proposed perceptual choice circuits for suboptimal (left) and optimal (right) computations. Orientation tuned neurons converge onto the decoder with excitatory weights biased towards target-preferring neurons. We propose that mice adopt a suboptimal computation implemented in a feed-forward excitatory circuit that lacks lateral inhibition from distractor-preferring neurons.

To test this hypothesis, we generated an optimal task-specific decoder by training a logistic regression in the control condition to correctly discriminate distractor and target stimuli and then tested in the two adaptation conditions. Indeed, the weights found by the logistic regression were also not equally distributed across the population. As expected, neurons preferring target orientations tended to have positive weights (p<10^−5^; Student’s t-test; **Figure 7b**), while neurons preferring stimuli closer to the distractor orientation tended to have negative weights (p<0.05). However, the logistic regression also failed to correctly predict the direction of the behavioral effects (auROC-22.5°: p=0.06; 0°: p=0.23; **Figure 7 – figure supplement 1b**). The same was true when we trained the decoder on all adaptation states.

This suggests that the behavior may be generated by properly, or at least positively, weighting the target neurons while largely ignoring the neurons that are tuned to the distractor. To test whether this positive weighting of the target responsive neurons is sufficient to explain the direction of the effects of adaptation on behavior, we set the weights of each neuron to one, zero or negative one according to its orientation preference. Indeed, if we generated a suboptimal weighting of inputs, where those neurons with preferences less than 30° were set to zero, while those above 30° were set to one, the decoder reliably predicted an increase in discrimination threshold and a decrease in FA rate with adaptation (auROC for 250 vs 750 ms: 22.5°: p<0.01; 0°: p<0.05; paired t-test; **Figure 7c**). The effect is not significant, though trends in the same direction, if the threshold for positive weights is set at 60° (22.5°: p=0.15; 0°: p=0.25; paired t-test). However, any addition of negative weights to distractor preferring neurons inverted the relationship between adaptation states such that increasing adaptation predicted a decrease in threshold and no change in false alarm rate (<15°: auROC-22.5°: p<0.01; 0°: p=0.25; <30°: auROC-22.5°: p<0.05; 0°: p=0.19; paired t-test; **Figure 7c**). Thus, the observed effects of adaptation on orientation discrimination are consistent with a suboptimal computation in which the downstream perceptual choice circuit performs an all-positive integration of the population activity (**Figure 7d**).

## Discussion

In order to determine how decision-making circuits use information from orientation tuned neuronal populations, we trained the mice to perform a multi-interval, go/no-go orientation discrimination task. We find that adaptation impairs the animal’s ability to discriminate target orientations both in terms of threshold and fraction of rewarded trials (22.5° target, accounting for difference in false alarm rate: 250 ms-24% trials rewarded; 750 ms: 34.7% trials rewarded). These behavioral data are consistent with a suboptimal computation in which the perceptual choice circuit relies on neurons that prefer target stimuli but fails to appropriately negatively weight neurons that prefer the distractor. This suggests that the brain is not always optimal, and may instead be more opportunistic in its approach for solving perceptual tasks.

In this study we use the term optimal to describe the computation purely from the perspective of the best possible use of the information present in the network that can be extracted linearly. Indeed, the optimal computation (using the neuronal weights from the logistic regression) does significantly better on our discrimination task than the suboptimal computation (using the neuronal weights from the behavioral fit) that discards the activity of distractor-preferring neurons (percent correct-optimal: 97%; suboptimal: 90%; p<10^−6^). Thus, the finding that the brain favors a suboptimal computation to discriminate orientation may tell us something about the constraints of the circuit. All of the optimal computations that we considered, including orientation estimation and the task-specific logistic regression, require the negative weighting of specific populations. Such a negative weighting cannot be achieved mono-synaptically and requires a neuron-specific recruitment of inhibition and so may be more difficult to learn (**Figure 7d**).

On the other hand, the computation suggested by our neural fit to the behavioral data relies only on an excitatory, feed-forward circuit. The perceptual choice circuit may learn though experience to rely on the activity of neurons which increase their activity in response to targets. This computation is amenable to a simple associative learning rule (Law and Gold, 2009; Znamenskiy and Zador, 2013; Xiong et al., 2015). For instance, the positive weighting of target neurons might be achieved through Hebbian long-term synaptic potentiation, while the lack of weight on the distractor neurons might be achieved through long-term depression. This proposed circuit could be directly tested by new tools that allow for the specific activation of functional subsets of neurons in behaving animals (Mardinly et al., 2018).

The demonstration that the mice are using a suboptimal computation is surprising, though there is certainly precedence for this (Beck et al., 2012; Ho et al., 2012; Oh et al., 2016). It is possible that the strategy used to train the mice to perform this task, or the specific task parameters used, supported the development of this suboptimal computation. For instance, the optimal circuit imposes at least one synaptic delay and requires additional integration time in the recurrent network. The suboptimal computation may thus support the fast decision-making needed in the discrimination task. This may be a general approach for fast decision-making: there is evidence that the human cortex uses suboptimal computations that ignore distractors or weakly informative features, especially under time constraint (Ho et al., 2012; Oh et al., 2016).

Notably there are multiple computations, general and task-specific, that can be used to successfully solve our discrimination task. The effects of adaptation (**Figure 4**) and subtle changes in distractor orientation on the animals’ performance (**Figure 6**) are consistent with the mice having adopted task-specific strategies for both target identification and discrimination. The finding that the mice use task-specific strategies is not surprising. Evidence from human behavioral experiments suggest that participants are likely to adopt task-specific strategies when they are trained on a small number of conditions (Fulvio et al., 2014) and experience large numbers of trials (Ramachandran and Braddick, 1973; Ball and Sekuler, 1982; Zhang et al., 2010).

While the task-specific, suboptimal perceptual choice circuit is viable, it leaves the animal vulnerable to systematic error. By relying on the absolute firing rate of a subset of neurons, anything that increases firing rates can be mistaken for a target. We think that this is why the mice have a high FA rate following long intervals: the recovery from adaptation with long intervals results in larger than expected firing rates, making the animal respond as if a target had been presented (**Figure 4d-g**). Indeed, reducing the firing rates in V1 by optogenetic activation of inhibitory neurons results in a decrease in hit and FA rates, consistent with the hypothesis that the perceptual choice circuit is summing the total activity in this area (**Figure 5f**). Moreover, the disproportional decrease in FA rates (and increase in threshold) with suppression of V1 is consistent with effects of ISI on behavior acting through effects on sensory coding, and not through more cognitive mechanisms like forward masking and attentional blink (Raymond et al., 1992; Macknik and Livingstone, 1998; Alwis et al., 2016). Notably, these phenomena also tend to act on much shorter time-scales (tens of milliseconds) than intervals used in this study, making them unlikely candidates to explain the effects of ISI on behavior. Finally, the lack of effect of ISI on lapse rate also argues against a cognitive mechanism for the effects of ISI on behavior.

The neuronal data that we used to generate the model predictions was collected from naïve mice that were passively viewing the visual stimuli. This was done to generate full orientation tuning curves in the adapted and unadapted conditions, as well as to avoid contamination of non-sensory signals. However, this means that the effects of training or active behavioral engagement are not included in our model predictions. Since the tuning of visual cortical neurons can be affected by visual experience (Schoups et al., 2001; Kreile et al., 2011; Goltstein et al., 2013), it is possible that our task training paradigm (abundance of 0° distractors and 90° targets, 6 day/week, >3 months) induced a change in the representation of orientation in V1 neurons. However, the orientation identification models are designed to account for skewed distributions, and do an excellent job of predicting the orientation of the stimulus despite the over-abundance of cardinal orientation preferring neurons (**Figure 3b**). Thus, we do not expect that experience-dependent changes in the representation of orientation would substantially affect orientation identification models’ predictions. On the other hand, the degree of adaptation is dependent on both the number of adapters preceding the target (**Figure 4 - figure supplement 1g-h**) as well as task engagement (Keller et al., 2017), and therefore the mapping of the neural data onto the behavioral data may not be straightforward. Nonetheless, we were able to reliably fit the neural data to the behavioral data (**Figure 7 – figure supplement 1a**), suggesting that there may in fact be a linear transform of firing rates across behavioral state.

Our behavioral, physiological and computational approaches reveal that the circuit adopts a suboptimal computation to solve an orientation discrimination task. This reveals that models for decoding sensory signals must be rigorously tested with experiments, and that the perceptual choice circuit may not always be optimized for the best use of available information. In this case, while ignoring important information leaves the mouse vulnerable to the effects of adaptation, it enables both fast associative learning and fast decision making via a simple feedforward circuit.

## Methods

### Animals

All animal procedures conformed to standards set forth by the NIH, and were approved by the IACUC at Duke University. 33 mice (both sexes; 3-24 months old; singly and group housed (1-4 in a cage) under a regular 12-h light/dark cycle; C57/B6J (Jackson Labs #000664) was the primary background with up to 50% CBA/CaJ (Jackson Labs #000654)) were used in this study. Ai93 (*tm93.1(tetO-GCaMP6f)Hze*, Jackson Labs #024103; n = 4) and Ai94 (*tm94.1(tetO-GCaMP6s)Hze*, Jackson Labs #024104; n = 8) were crossed to *EMX1-IRES-Cre (Jackson Labs #005628)* and *CaMK2a-tTA (Jackson Labs #003010)* to enable constitutive GCaMP6 expression for *in vivo* imaging experiments. *Pvalb-cre* (*tm1(cre)Arbr*, Jackson Labs #008069; n = 13), *VGAT-ChR2-EYFP* (*Slc32a1-COP4*H134R/EYFP*, Jackson Labs #014548; n= 3) and Emx1-IRES-Cre (*tm1(cre)Krj*, Jackson Labs # 005628; n = 2) were crossed to C57/B6J mice for *in vivo* extracellular electrophysiology (n = 4) and behavior (n = 14) experiments. *Gad2-IRES-cre* (Gad2^tm2(cre)Zjh^, Jackson Labs #010802; n = 2) and C57/B6J (n = 1) mice were crossed to CBA/CaJ for eye-tracking experiments.

### Cranial window implant

Dexamethasone (3.2 mg/kg, s.c.) and Meloxicam (2.5 mg/kg, s.c.) were administered at least 2 h before surgery. Animals were anesthetized with ketamine (200 mg/kg, i.p.), xylazine (30 mg/kg, i.p.) and isoflurane (1.2-2% in 100% O_2_). Using aseptic technique, a headpost was secured using cyanoacrylate glue and C&B Metabond (Parkell), and a 5 mm craniotomy was made over the left hemisphere (center: 2.8 mm lateral, 0.5 mm anterior to lambda) allowing implantation of a glass window (an 8-mm coverslip bonded to two 5-mm coverslips (Warner no. 1) with refractive index-matched adhesive (Norland no. 71)) using Metabond.

The mice were allowed to recover for one week before habituation to head restraint. Habituation to head restraint increased in duration from 15 min to >2 h over 1-2 weeks. During habituation, imaging and electrophysiology sessions, mice were head restrained while allowed to freely run on a circular disc (InnoWheel, VWR). Wheel revolutions were monitored with a digital encoder.

### Visual stimulation

Visual stimuli were presented on a 144-Hz LCD monitor (Asus) calibrated with an i1 Display Pro (X-rite). The monitor was positioned 21 cm from the contralateral eye. Circular 30° gabor patches containing static sine-wave gratings (0.1 cycles per degree) alternated with periods of uniform mean luminance (60 cd/m^2^). Visual stimuli for imaging, electrophysiology and behavior experiments were controlled with MWorks (http://mworks-project.org).

Three visual stimulus protocols were used for imaging experiments: 1) Paired-pulse, same orientation (**Figure 1**); 2) Paired pulse, different orientation (**Figure 2**); and 3) Six-pulse, random interval, random target (**Figure 4 – figure supplement 1g**). In protocol 1 (n = 5 mice), two static, 100 ms sine-wave gratings of the same orientation (0°, 30°, 60°, 90°, 120°, or 150°) were successively presented with a variable inter-stimulus interval (ISI: 0.25, 0.5, 1, 2, or 4 s) and an inter-trial interval (ITI) of 4s. Measurement of adaptation was averaged across all orientations, except in **Figure 1d**. In protocol 2 (n = 12 mice), a static, 100 ms sine-wave vertical grating (0°; “adapter”) was followed by a 100 ms grating (“test”) of varying orientation (0°, 22.5°, 45°, 67.5°, 90°, 112.5°, 135°, or 157.5°) after a variable ISI (250 or 750 ms), with an ITI of 8 s. On 30% of trials, the first stimulus was omitted to measure the non-adapted (control) tuning curve. In protocol 3 (n = 3 mice), five static, 100 ms vertical sine-wave gratings were successively presented followed by a 100 ms grating of varying orientation (30° or 90°), with an ITI of 8 s. ISIs (250, 500 or 750 ms) were selected on a presentation-by-presentation basis.

For electrophysiology experiments only protocols 1 (n=4 mice) and 3 (n=4 mice) were used. Stimuli were the same as in the imaging protocols except 1) only one orientation (0°) was used in protocol 1, and the ITI was 10s; and 2) the targets in protocol 3 were 22.5° and 90°, and the number of baseline presentations preceding the target was also randomized from 2-9 with ITI of 4 s. For all stimulus protocols, all orientations and interval conditions were randomly interleaved.

### Retinotopic mapping

Retinotopic maps generated from either intrinsic autofluorescence or GCaMP signals. For intrinsic autofluorescence imaging, the brain was illuminated with blue light (473 nm LED (Thorlabs) or 462 ± 15 nm band filter (Edmund Optics)), and emitted light was measured through a green and red filter (500 nm longpass). Images were collected using a CCD camera (Rolera EMC-2, Qimaging) at 2 Hz through a 5× air immersion objective (0.14 numerical aperture (NA), Mitutoyo), using Micromanager acquisition software (NIH). Stimuli were presented at 4-6 positions (drifting, sinusoidal gratings at 2 Hz) for 10 s, with 10 s of mean luminance preceding each trial. Images were analyzed in ImageJ (NIH) to measure changes in fluorescence (dF/F; with F being the average of all frames) to identify primary visual cortex (V1) and the higher visual areas. GCaMP imaging followed an identical procedure except light was collected with a bandpass filter (520 ± 18 nm) and total trial duration was reduced to 10 s. Vascular landmarks were used to identify targeted sites for imaging, electrophysiology and optogenetics experiments.

### Viral injection

We targeted V1 in *Pvalb-cre* mice (n=2) for expression of Channelrhodopsin2 (ChR2). Dexamethasone (3.2 mg/kg, s.c.) was administered at least 2 h before surgery and animals were anesthetized with isoflurane (1.2-2% in 100% O_2_). The coverslip was sterilized with 70% ethanol and the cranial window removed. A glass micropipette was filled with virus (AAV5.EF1.dFloxed.hChR2.YFP (UPenn CS0384)), mounted on a Hamilton syringe, and lowered into the brain. 50 nL of virus were injected at 250 and 500 µm below the pia (30 nL/min); the pipette was left in the brain for an additional 10 minutes to allow the virus to infuse into the tissue. Following injection, a new coverslip was sealed in place, and an optical cannula (400 µm diameter; Doric Lenses) was attached to the cranial window above the injection site. Optogenetic behavioral experiments were conducted at least two weeks following injection to allow for sufficient expression.

### Two-photon calcium imaging

Images were collected with a two-photon microscope controlled by Scanbox acquisition software (Neurolabware). Excitation light (920 nm) from a Mai Tai eHP DeepSee laser (Newport) was directed into a modulator (Conoptics) and raster scanned onto the brain with a resonant galvanometer (8 kHz, Cambridge Technology) through a 16X (0.8 NA, Nikon) or 25X (1.05 NA, Nikon) water immersion lens. Average power at the surface of the brain was 30-50 mW. Frames were collected at 30 Hz (256 lines) for a field of view of ~700 × 400 μm on a side. Emitted photons were directed through a green filter (510 ± 42 nm band filter (Semrock)) onto GaAsP photomultipliers (H10770B-40, Hamamatsu). Images were captured at a plane 207 ± 4 μm below the pia (range 180-250 μm). Frame signals from the scan mirrors were used to trigger visual stimulus presentation for reliable alignment with collection.

### Eye-tracking

Images were collected at 30 Hz with a Genie Nano CMOS camera (Teledyne Dalsa) using a 695 nm LP filter (Midopt) controlled by Scanbox acquisition software. IR illumination (920 nm) was provided from the two-photon laser through the cranial window.

### Extracellular electrophysiology

Electrophysiological signals were acquired with a 32-site polytrode acute probe (either A4×8-5mm-100-400-177-A32 (4 shanks, 8 site/shank at 100 μm spacing) or A1×32-Poly2-5mm-50s-177-A32, (1 shank, 32 sites, 25 μm spacing), NeuroNexus) through an A32-OM32 adaptor connected to a Cereplex digital headstage (Blackrock Microsystems). Unfiltered signals were digitized at 30 kHz at the headstage and recorded by a Cerebus multichannel data acquisition system (Blackrock Microsystems). Visual stimulation synchronization signals were also acquired through the same system via a photodiode directly monitoring LCD output.

On the day of recording, the cranial window was removed, and a small durotomy performed to allow insertion of the electrode into V1. A ground wire was connected via a gold pin cemented in a burrhole in the anterior portion of the brain. The probe was slowly lowered into the brain (over the course of 15 min with travel length of around 800 μm) until the most superficial recording site was in the brain and allowed to stabilize for 45 - 60 min before beginning recordings. A fluorescent dye (diI, Life Technologies) was painted on the back of the probe prior to recording and the probe position was thus confirmed post hoc in histological sections.

### Behavioral task

Animals were water scheduled and trained to discriminate orientation by manipulating a lever. The behavior training and testing occurred during the light cycle. We first trained mice to detect full-field, 90° orientation changes from a static grating. Most mice (n=12) were trained with a 0° distractor; however, 2 mice were trained with a 45° distractor. On the initial days of training, mice were rewarded for holding the lever for at least 400 ms (required hold time) but no more than 20 s (maximum hold time). At the end of the required hold time, the target grating appeared and remained until the mouse released the lever (or the maximum hold time expired). Typically, within two weeks of training, the mice began releasing the lever as soon as the target appeared. Once the animals began reliably responding to the target stimulus, we added a random delay between lever press and the target presentation to discourage adoption of a timing strategy. Over the course of the next few weeks, the task was made harder by (in roughly chronological order): 1) increasing the random delay, 2) decreasing the target stimulus duration and reaction time window, 3) removing the stimulus during the ITI, 4) shrinking and moving the stimuli to more eccentric positions, 5) adding a mean-luminance ISI to mask the motion signal in the transition from distractor to target, and finally 6) reducing the difference between the distractor and the target. Delays after errors were also added to discourage lapses and early releases.

In the final form of the task, each trial was initiated when the ITI had elapsed and the mouse had pressed the lever. Trial start triggered the presentation of a 100 ms static sinusoidal, gabor patch (30° in diameter, positioned at an eccentricity of 30° - 40° in azimuth and 0° - 10° in elevation) followed by an ISI randomly selected on a presentation-by-presentation basis (250, 500 or 750 ms). For a subset of mice (n=3), the ISI was fixed for a given trial but randomly interleaved on trial-by-trial basis (250 or 500 ms). The target appeared with a random delay (flat distribution) after the first two presentations on each trial and was randomly selected from a fixed set of values around each animal’s threshold. Mice received water reward if they released the lever within 100-650 ms (sometimes extended to 1000 ms) after a target occurred. However, for calculating hit and false alarm (FA) rate (**Figures 4, 5 and 6**), we use a narrower reaction window (200-550 ms) to ensure that the majority of the releases in this window are due to stimulus driven responses and have independent reaction windows for adjacent stimuli with short ISIs. Mice were initially trained with a constant distractor stimulus (**Figure 4 and 5**) and were later tested with randomly interleaved distractor orientations (the trained orientation and stimuli −15° and +15° from the trained orientation) selected on a trial-by-trial basis (**Figure 6**).

Of the 11 mice presented in **Figure 4**, some (n=5) were initially trained with a single (250 ms) ISI, whereas others (n=6) were immediately introduced to having the fully interleaved condition (presentation-by-presentation selection of one of three ISIs). Training history had no significant effect on the relationship between ISI and threshold (two-way anova; main effect of ISI (p<0.005); main effect of training (p=0.41)). Notably, the incorporation of targets close to the distractor occurs during the final stages of training; thus, the mice learn relatively late in training that the targets lie within only one quadrant of orientation space. Nonetheless, the FA rates from the variable baseline task (**Figure 6**) reveal that the mice understand this contingency: if the mice continued to use the strategy learned when there was only a 90° target, then the 15° and - 15° distractors should have similar FA rates.

For optogenetic stimulation (**Figure 5**), we delivered blue light to the brain though the cannula from a 473 nm LED (Thorlabs) or a 450 nm laser (Optoengine) and calibrated the total light intensity at the entrance to the cannula (0.1 - 0.3 mW). On randomly interleaved trials, the light was turned on at the time of lever press and remained on until lever release. Behavioral control was done with MWorks, and custom software in MATLAB (MathWorks) and Python (http://enthought/com).

## Data processing

### Image processing and analysis

All image processing and analysis was performed in MATLAB. The image stack was registered to a stable, average field of view using sub-pixel registration methods to correct for motion along the imaged plane (*x-y* motion). For segmentation of visually driven neurons, we used semi-automated segmentation algorithms to select regions of interest (cell masks) from the average change in fluorescence (*dF/F*, where *F* is the average fluorescence in the 20 frames (~660 ms) preceding the first stimulus in each trial) evoked in response to each stimulus type. Fluorescence time courses were generated by averaging all pixels in a cell mask. Neuropil signals were removed by first selecting a shell around each neuron (excluding neighboring neurons), estimating the neuropil scaling factor (by maximizing the skew of the resulting subtraction), and removing this component from each cell’s time course.

Visually evoked responses were measured as the difference in the *dF/F* before (baseline window: average of three frames (~100 ms) around stimulus onset) and during the response (response window: average of three frames around the peak response; window was selected separately for each experiment to account for variability in response latencies and indicator kinetics). “Responsive cells” were chosen as having statistically significant visually evoked responses to at least one of the stimulus types (bonferroni-corrected paired t-test) or all stimuli (paired t-test), and the maximum derivative in the *dF/F* occurred before the end of the response window (to eliminate cells strongly driven by the removal of the stimulus). Using these criteria, 245/279, 473/587, and 202/239 cells were included for visual stimulation protocols 1-3, respectively. All measurements are the average of at least 7 trials of the same type.

Cells imaged in protocol 2 (paired-pulse, different orientation) were further selected based on the reliability of their orientation tuning. Responses in the control (no adapter) condition were fit with a Von Mises function:

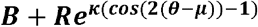

where *B* is the baseline firing rate, *R* is the modulation rate, κ is the concentration, and μ is the preferred orientation. To measure the reliability of the fit, the fit was repeated 1000 times resampling trials with replacement. Only cells for which 90% of the bootstrapped fits were within 22.5° of the original fit were included in further analysis (241/473 cells).

Analysis of the effects of adaptation on tuning (e.g. the preferred orientation and the orientation selectivity index (OSI); **Figure 2e-g**) were derived from the von Mises fit to the data. OSI was measured as:

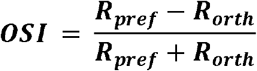

where *R*_*pref*_ is the cell’s response at the preferred orientation (Pref: maximum of the fit) and *R*_*orth*_ is the response to the orthogonal orientation. In the case that *R*_*orth*_ was negative, it was set to 0. Cells were grouped according to the distance of their preferred orientation from the adapter (cells that prefer 22.5° and 157.5° were in the same group; **Figure 2e-g**).

For protocols 1 and 3, each cell’s responses were normalized to the average response to the first stimulus. For protocol 2, each cell’s responses were normalized to *R*_*pref*_.

### Electrophysiology processing and analysis

Individual single units were isolated using the SpyKing CIRCUS package (http://spyking-circus.readthedocs.io/en/latest/). Raw data were first high pass filtered (> 500 Hz) and spikes were detected when a filtered voltage trace crossed threshold (9-13 median absolute deviations computed on each channel). A combination of density-based clustering and template matching algorithms were used to automatically cluster the spikes. The resulting clusters were then inspected and adjusted manually using a MATLAB GUI. Clusters with refractory period violations (< 2 ms, >1% violation) in the auto-correlogram and that were not stable across the whole recording session were discarded from the dataset. Clusters were combined if they met each of three criteria by inspection: 1) similar waveforms; 2) coordinated refractory periods in the cross-correlogram; 3) similar inter-spike interval distribution shape. Unit position with respect to the recording sites was calculated as the average of all site positions weighted by the waveform amplitude of each site. All the subsequent analysis was performed in MATLAB.

Local field potential (LFP) and current source density analysis (CSD). For LFP recording, the extracellular raw signal was band pass filtered from 1 to 200 Hz and downsampled to 10 kHz. The CSD was computed from the average LFP by taking the discrete second derivative across the electrode sites that were linearly spaced across cortical depth, and interpolated to produce a smoothed visually driven CSD profile. This analysis transformed the LFP signal into the locations of current sources and sinks, revealing a patterned laminar distribution of sinks in V1 after the visual onset: an initial sink in layer 4 (latency: ~50 ms), followed by a sink in layer 2/3 and finally a weak and sustained sink in layer 5. Therefore, guided by the visually-evoked CSD map, retrospective histology, and relative depth of recordings relative to the pia surface, layer 2/3 units were identified and chosen for comparison with the two-photon imaging dataset.

Visually-evoked responses of each unit in layer 2/3 of V1 were measured based on average peri-stimulus time histograms (PSTHs, bin size: 20 ms) over repeated presentations (>25 trials) of the same stimulus. Response amplitudes were measured on a trial-by-trial basis: by subtracting the firing rate at the time of the visual stimulus onset from the value at the peak of the average PSTH within a window of 0-100 ms after the visual onset. “Responsive cells” were chosen as having statistically significant visually-evoked responses (first baseline response, averaged over 0-100 ms before the visual onset, vs visually-evoked responses, averaged over 0-100 ms after the visual onset; paired t-test; this analysis window excluded off-responsive units from analysis). For protocol 1, we only included the responsive units that had no significant difference in response to the first stimulus (the adapter) across five ISIs (one-way ANOVA). Similarly, for protocol 3, we included responsive units for the analysis of the normalized firing rate binned by cycle numbers (**Figure 4 - figure supplement 1h**). Using these criteria, 30/39 and 17/25 layer 2/3 cells were included for visual stimulation protocols 1 and 3, respectively.

### Behavior processing and analysis

All behavioral processing and analysis were performed in MATLAB. All trials were categorized as either an early release, hit, or miss based on the time of release relative to target onset: responses occurring earlier than 100 ms after the target were considered early releases; responses occurring between 200 and 550 ms after a target were considered hits; failures to respond before 550 ms after the target were considered misses. Behavioral sessions were manually cropped to include only stable periods of performance and were further selected based on the following criteria: 1) at least 50% of trials were hits; and 2) less than 35% of trials were early releases. Based on these criteria, the data in **Figure 4** included 17± 3 sessions (range: 5-46) for each mouse with an average of 6348 ± 815 trials per mouse (range: 2593- 11857); the data in **Figure 5** included 24 ± 5 (range: 11-35) sessions for each mouse with 7524 ± 1582 trials (range: 5021-11710), respectively; and the data in **Figure 6** included 20 ± 5 sessions (range: 6-40) for each mouse with an average of 4904 ± 941 trials per mouse (range: 2284 - 8236).

Hit rate was computed from the number of hits and misses for each stimulus type:

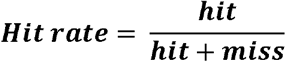

Lapse rates were measured as (1-Hit rate) for 90° targets. Most mice had low lapse rates (11/11 mice were below 10%) for the task in **Figure 4**. However, as mice age their reaction times become slower, thereby inflating the lapse rate; we think that this effect explains the increased lapse rate during optogenetic suppression of V1 (only 1/4 mice were below 10% in **Figure 5**).

Hit rates across stimulus types were fit with a Weibull function to determine discrimination thresholds (50% of the upper asymptote to account for lapse rate). No correction was made for FA rate. Threshold CIs were estimated via nonparametric bootstrap resampling trials with replacement.

All distractor stimulus presentations were categorized as either a CR or a FA: responses occurring between 200 and 550 ms after a distractor stimulus were considered FAs; presentations where the mouse held the lever for at least 550 ms after the distractor stimulus were considered CRs. FA rate was computed from the total number of FAs and CRs in the session:

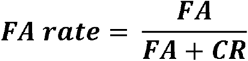

Signal detection theory (Green and Swets, 1966) was applied to calculate sensitivity (d’) and bias (c) given the measured hit and FA rate as follows:

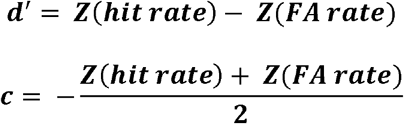

where Z is the inverse of the cumulative distribution function of the normal Gaussian distribution.

For matching the trial length across ISIs (Figure 4 – figure supplement 1d), trial length was first binned every 0.5 s within the range of 1.2 - 6.2 s (for hits and misses) and 1.2 - 4.2 s (for FAs and CRs) respectively. Within each bin, the same number of trials were chosen for all ISIs to ensure the average trial length was not significantly different. The selected trial number for each bin was determined by the minimum number of trials across ISIs in that bin for each mouse.

For calculating fraction of rewarded trials (P_rew_) given the 22.5°- Hit and FA rates of each ISI (250 and 750 ms), we first simulated 1000 trials where the 22.5° target is presented after six distractor 0°presentations. For a known FA rate, we calculated the fraction of early trials (P_early_) that were unrewarded assuming every 1/FA number of distractor presentations would generate an early release. Then the P_rew_ was calculated as follow:

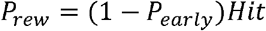

### Eye-tracking processing and analysis

All eye-tracking data was analyzed in MATLAB. The size of the pupil on each frame was extracted using the built-in function imfindcircles. Pupil size was normalized to the maximum radius measured during the session and quantified as the average measured in a 250 ms window preceding the target stimulus.

### Modeling

For all modeled decoders, neurons imaged in different mice were analyzed separately and only datasets with at least 10 well-fit neurons (using the criteria above) were included in these analyses (10/12 experiments). As a result, error bars in **Figures 3**, **7** and associated supplements represent standard error of the mean across data sets.

Log likelihood functions were generated using two methods: The first followed the equation(Jazayeri and Movshon, 2006):

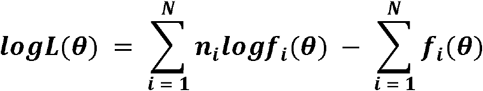

where *N* is the number of neurons in the population, f_i_*(Θ)* is each neuron’s normalized tuning function and *n*_*i*_ is each neuron’s response to a given stimulus. The tuning functions were obtained from well-fit, responsive neurons, in the non-adapted condition (under the assumption that the decoder is unaware of the effects of adaptation(Seriès et al., 2009)), and were used to predict the likelihood of the stimulus given single trial responses (*n*_*i*_) that varied with adaptation.

Because our neuronal data does not adhere to the assumptions that spiking is Poisson and independent, we also used a second more empirical method based upon the assumption that our population of neurons represented von Mises over orientation using a linear PPC(Ma et al., 2006). This approach assumes that the likelihood function takes the form:

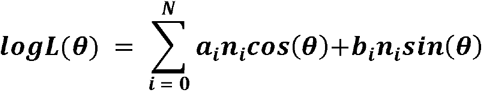

where **n**_0_ is assumed to be 1 and the *a*_i_’s and *b*_i_’s are discovered by maximizing the likelihood of the empirically observed joint distribution of the presented stimulus and the neural response. This results in a convex optimization problem which we solved using gradient ascent. Because this method is prone to overfitting resulting in poor cross-validation performance when the number of units greatly exceeds the number of trials per condition, in datasets with more than 15 well-fit neurons, we used only the 15 best-fit neurons (as measured by their 90% CI). As a control, we also preprocessed our neural data using principle component analysis and eliminated all but the 15 dominant modes of variability and refit the empirically generated log likelihood. This had no effect on our results. The number 15 was chosen to optimize performance on the cross validated data set, however, we note that choosing values between 12 and 16 did not substantially change our results.

The single-trial log likelihood functions were then used to determine either the MAP estimate (Method 1: **Figure 3e**; Method 2: **Figure 3b-c**) or the posterior probability distribution (“optimal decoder”: **Figure 3d**). We also used a standard population vector decoder to estimate the orientation (**Figure 3f**). For this decoder a preferred stimulus value, *θ*_i_, was extracted from a parameterized fit to the tuning curve of each well-fit unit. Estimated orientation for each trial was then obtained from the equation:

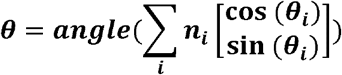

Logistic regression was performed using MATLAB’s glmfit routine with a binary observation set to be 1 when the presented stimulus was not vertical (the adapter orientation).

Estimate biases were computed by taking the mean estimate as a function of stimulus and adaptation condition. Because there was no clear trend in estimate standard deviation as a function of presented orientation, we computed estimate variance in each adaptation condition by removing the bias associated with each stimulus value and concatenated the resulting estimate residuals into a single vector. In order to account for differences in the information content of each data set, estimate variances (and standard deviations) were normalized by the variance of the control condition of each data set (35±14 degrees). We then concatenated the resulting normalized residuals from each data set into a single data set to measure the mean and associated standard error of the residual variances; these measures are shown in the inset of **Figure 3b**.

To compute the auROC we treated our estimates of orientation (or in the case of the optimal decoder and logistic regression: the probability that the orientation was not the adapter orientation) as decision variables and computed false positives and correct detections for 400 uniformly sampled values of the decision criterion chosen to span all observed values of the decision variable. For the sum decoder we simply treated the total population activity on each trial as the decision variable. The auROC was then computed numerically using the trapezoid rule.

Note that in all cases we report only cross-validated results (using leave one out cross-validation in the control condition). This is because parameters of our neural decoders were all fit using only the control condition and not our two adaptation conditions. The only exception to this is for the calculation of neuronal weights using the logistic regression; in this case, to make it comparable to the estimation of neuronal weights from the fit to the behavior, we trained the decoder on the 750 and 250 ms conditions. The results for all decoders were largely unchanged if we obtained parameters from either the 750 ms condition or all conditions simultaneously and cross-validate using a leave one out procedure. However, we note that logistic regression weights are not sensitive to this choice of training data sets.

To determine the properties of a neural decoder capable of fitting our behavioral data we summarized the behavioral data in the two adapted conditions for the 11 animals used in **Figure 4**. Since different orientations were sampled for each of the mice, we performed a spline fit of each psychometric function and averaged across all animals (**Figure 7 – figure supplement 1a**). For each neuronal dataset we assumed that behavior was generated by sampling from a posterior distribution that had the form of a logistic regression, i.e.:

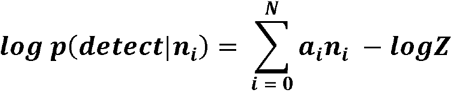

where *n*_0_ is once again assumed to be 1 and Z is the normalizer. The *a*_i_’s were then determined by minimizing the symmetrized Kullback-Leibler divergence between the average of this neurally predicted detection probability raised to some power and the behaviorally observed detection probability across stimulus and adaptation conditions. Once again, we used gradient descent to perform this optimization. These behaviorally generated weights strongly correlated with the weights discovered by applying logistic regression (correlation coefficient=0.54) and the resulting mean squared error between the neurally generated psychometric curves and the behaviorally observed ones was 2e-4.

In order to assess the degradation in performance that results from use of a suboptimal decoder we computed percent correct in the following way. First the weights obtained from the behavioral model and logistic regression model fit to the 250 and 750 conditions were used to generate a set of decision variables for each data set and stimulus condition. A task relevant measure of percent correct was computed from these values by mimicking the statistics of the behavioral task (i.e. distractors are 8 times more common than targets and target stimulus values are uniformly distributed) for a range of potential decision criteria. The optimal decision criterion was selected by determining which provided the maximum value of percent correct. This resulted in two values for optimal percent correct, one for the behavioral weights and one for the logistic regression weights for each data set. Average and standard deviations for these percent correct values were then computed across data sets.

### Statistical analysis

Data were tested for normality using a Lilliefors test. While all behavioral measures and auROC estimates were normally distributed, distributions of neuronal responses were not. Thus, in the case that two distributions were compared we used a t-test in behavioral measures and auROCs, and a Wilcoxon rank sum test for neuronal responses; however, since ANOVA and post hoc Tukey HSD tests have been shown to be robust to non-normality (Driscoll, 1996), these tests were used for all data. Sample sizes were not predetermined by statistical methods, but our sample sizes of the neurons and behavior animals are similar to other studies. Data collection and analysis were not performed blind to experimental conditions, but all visual presentation conditions in either calcium imaging, extracellular recording or behavior testing are randomized. Moreover, the strength and timecourse of adaptation on neuronal responses was measured using two methods (**Figure 1**) with data collected by two experimenters.

### Data and code availability

All relevant data and code will be made available via GitHub and Globus.

## Acknowledgements

We thank B. Gincley and J. Sims for assistance with behavioral training; K. Leonard, M. Fowler and J. Isaac for surgical assistance; Z. Xu for assistance with software development; C. Hass, G. Field, C. Hull, S. Lisberger, M. Scanziani, A. Wilson, and X. Yao, for helpful discussions and comments on the manuscript. This work was supported by an NIH Director’s New Innovator Award (DP2-EY025439), the Pew Biomedical Trusts, and the Alfred P. Sloan Foundation (L.L.G).

## Author Contributions

M.J. and L.L.G. designed the experiments. M.J. collected and analyzed the electrophysiology and behavior data. L.L.G. collected and, together with J.M.B., analyzed the calcium imaging data. M.J., J.M.B., and L.L.G wrote the manuscript.

---

**Figure 4 – figure supplement 1.**
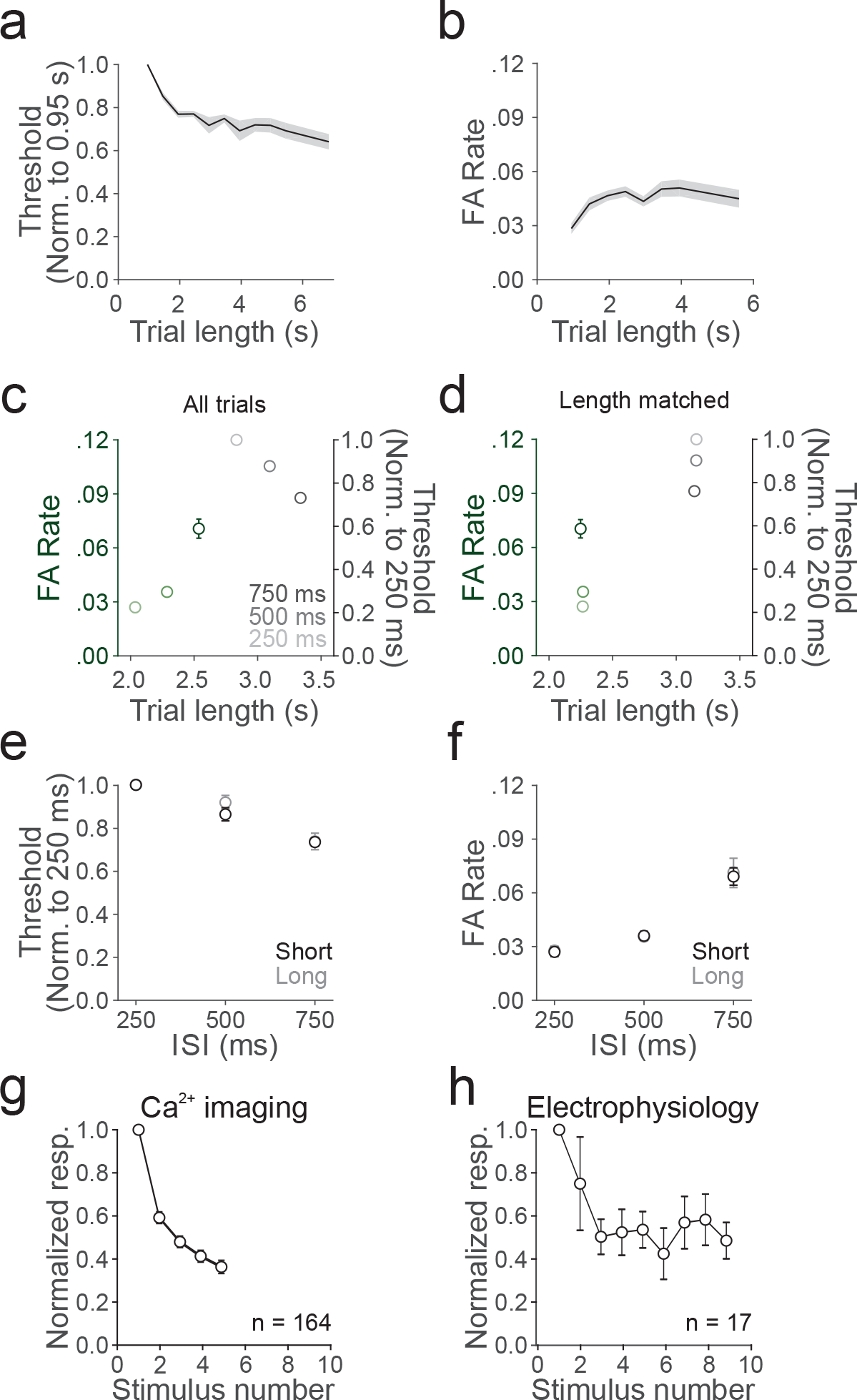
The effect of adaptation on behavior is independent of trial length. (**a**) Discrimination threshold as a function of trial length (bin size: 0.5 s). Values are normalized by the value at the first bin (center: 0.95 s). Shaded area is the SEM across 11 mice. (**b**) Same as **a**, for FA rate. (**c**) Summary of the change in threshold (normalized to 250 ms ISI, gray shades, right y-axis) and FA rate (non-normalized, green shades, left y-axis) as a function of average trial length. Note that by definition, the trial length for a FA is shorter than a hit. Error bars are SEM across 11 mice. (**d**) Same as **c**, but only including trials with matched length across ISIs. Note that there is no difference in average trial length across ISIs for discrimination threshold (p=0.83, one-way anova) and FA rate (p=0.24, one-way anova). (**e**) Summary of the change in threshold, normalized to the threshold in the 250 ms condition, as a function of ISI, for short (2-4 distractor presentations; black) or long (6-8 distractor presentations; gray) trials. (**f**) Same as **e**, for FA rate. (**g**) Normalized response amplitude to repeated presentations of distractors, as measured with Ca^2+^ imaging. Error is SEM across cells. (**h**) Same as **g**, measured with electrophysiology.

**Figure 4 – figure supplement 2.**
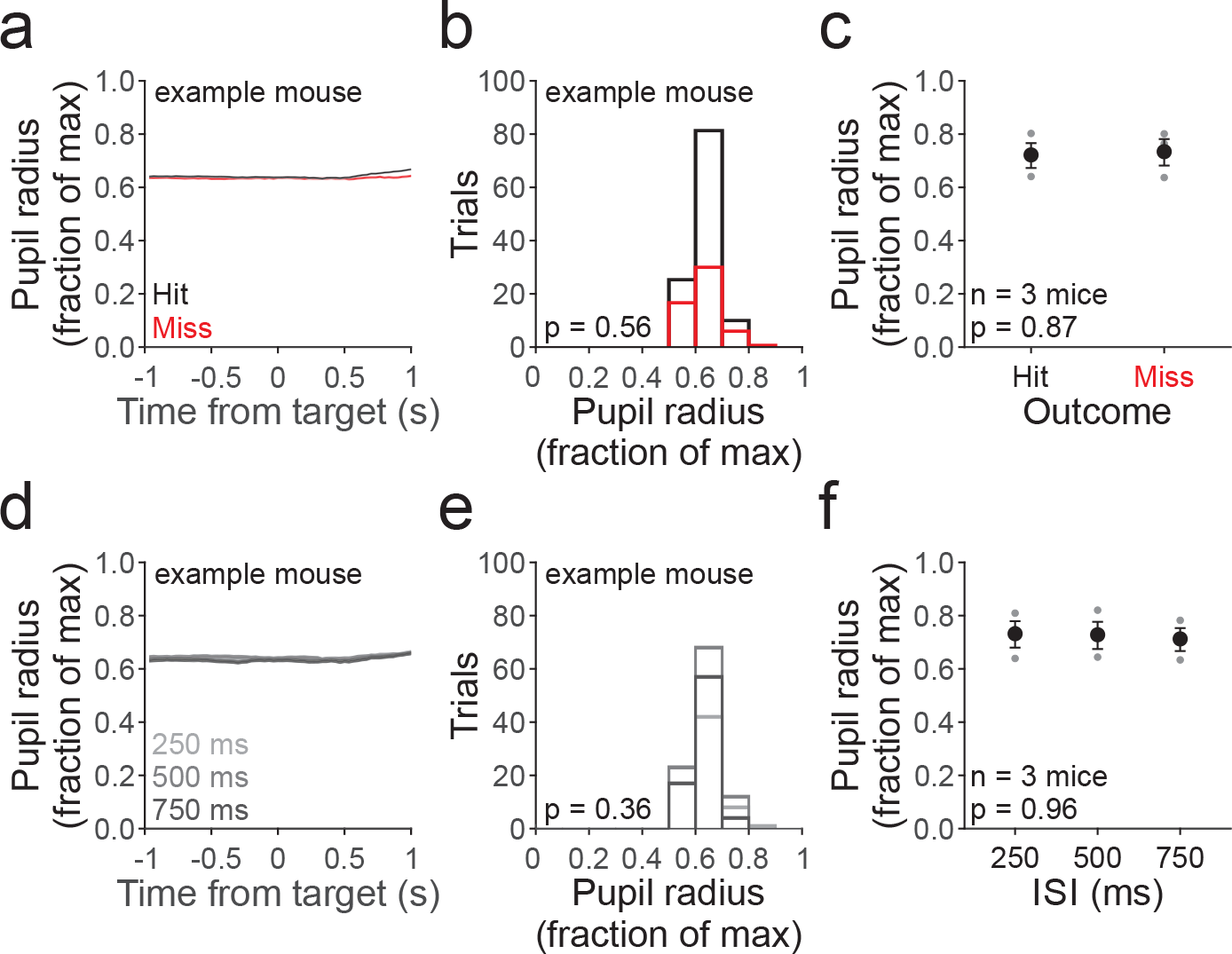
No relationship between pupil size and outcome or ISI. (**a**) Average time course of pupil radius for an example mouse normalized to the maximum radius measured during the experiment and aligned to time of stimulus target onset for hit (black) and miss (red) trials. Note the increase in pupil size after the target on hit trials, likely due to reward. Shaded error is +/− SEM across trials. (**b**) Histogram of average pupil size (in a 250 ms window preceding the target) for each hit (black) and miss (red) trial from same mouse as in a. P-value is from an unpaired t-test. (**c**) Summary of average normalized pupil radius for each mouse (gray circles) and across mice (black circles) by outcome. Errorbars are SEM across mice; p-value is from a paired t-test. (**d**) Average time course of normalized pupil radius divided by ISI. Same mouse and conventions as in **a**. (**e**) Histogram of average pupil size for trials divided by ISI; same conventions as in **b**. (**f**) Summary of average normalized pupil radius for each mouse (gray circles) and across mice (black circles) by ISI. Errorbars are SEM across mice; p-value is from a one-way anova.

**Figure 7 – figure supplement 1.**
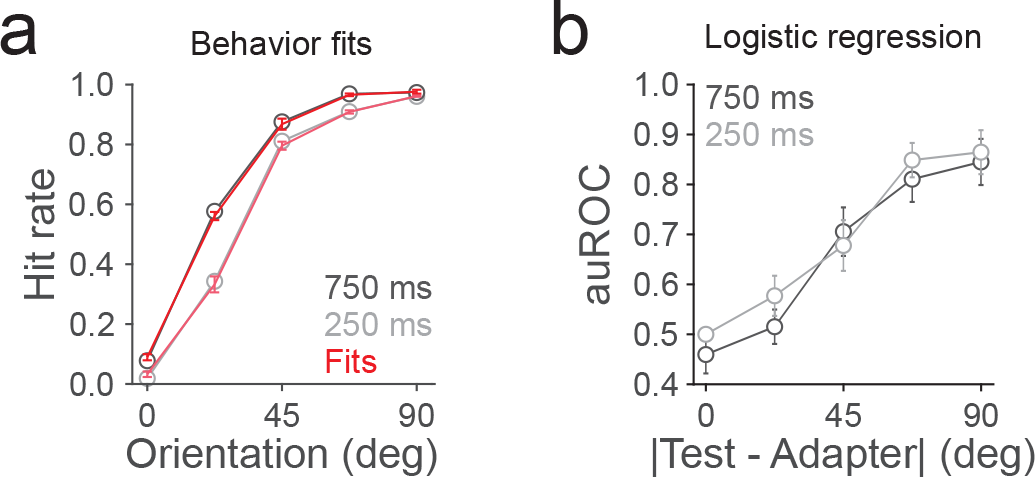
Logistic regression cannot account for effects of adaptation on behavior. (**a**) Fit (red) of the neural data to the behavioral data; hit rates are responses to targets (orientations > 0°) and distractors (0°) after 750 (dark gray) or 250 (light gray) ms recovery from adaptation. (**b**) Summary of the average auROC, found via logistic regression, as a function of stimulus distance from the adapter (n = 10 mice).

